# Investigating the Triple Code Model in Numerical Cognition Using Stereotactic Electroencephalography

**DOI:** 10.1101/2024.04.27.591485

**Authors:** Hao Tan, Alexander P. Rockhill, Christian G. Lopez Ramos, Caleb Nerison, Beck Shafie, Maryam N. Shahin, Adeline Fecker, Mostafa Ismail, Daniel R. Cleary, Kelly L. Collins, Ahmed M. Raslan

**Affiliations:** Departments of Neurological Surgery, Oregon Health & Science University, Portland, OR, 97239, USA

**Keywords:** numerical cognition, triple code model, numerical representation, intracranial EEG

## Abstract

The ability to conceptualize numerical quantities is an essential human trait. According to the “Triple Code Model” (TCM) in numerical cognition, distinct neural substrates encode the processing of visual, auditory, and nonsymbolic numerical representations. While our contemporary understanding of human number cognition has benefited greatly from advances in clinical imaging, limited studies have investigated the intracranial electrophysiological correlates of number processing. In this study, 13 subjects undergoing stereotactic electroencephalography for epilepsy participated in a number recognition task. Drawing upon postulates of the TCM, we presented subjects with numerical stimuli varying in representation type (symbolic vs. non-symbolic) and mode of stimuli delivery (visual vs. auditory). Time-frequency spectrograms were dimensionally reduced with principal component analysis and passed into a linear support vector machine classification algorithm to identify regions associated with number perception compared to inter-trial periods. Across representation formats, the highest classification accuracy was observed in the bilateral parietal lobes. Auditory (spoken and beeps) and visual (Arabic) number formats preferentially engaged the superior temporal cortices and the frontoparietal regions, respectively. The left parietal cortex was found to have the highest classification for number dots. Notably, the putamen exhibited robust classification accuracies in response to numerical stimuli. Analyses of spectral feature maps revealed that non-gamma frequency below 30 Hz held greater than chance classification value and could be potentially used to characterize format specific number representations. Taken together, our findings obtained from intracranial recordings provide further support and expand on the TCM model for numerical cognition.

## INTRODUCTION

The capacity to comprehend numerical quantities is remarkably phylogenetically conserved across several animal species (Brannon, 2010). However, the ability to deftly conceptualize and leverage abstract symbolic representations of numerical quantities is a uniquely human trait (Brannon, 2010, Sokolowski, 2017). Given the ubiquitous need to interact with quantities in daily life and the deleterious quality-of-life consequences associated with poor number literacy (Bynner & Parsons, 1997), developing further insights into the neural basis for our sophisticated numerical processing capabilities is of particular interest and value.

A leading framework for numerical cognition is the “Triple Code Model” (TMC) (Dehaene 1992, Dehaene & Cohen, 1995). The TCM postulates that cortical neuronal populations within the ventral temporal lobe, perisylvian region, and intraparietal sulcus are responsible for processing visual (“6”), auditory verbal (“six”), and nonsymbolic (“••••••”) numerical representation formats, respectively (Dehaene & Cohen, 1995). Studies utilizing functional magnetic resonance imaging (fMRI) and transcranial magnetic stimulation (TMS) have lent support to the TCM while also indicating a broader engagement of cortical regions beyond those originally proposed, such as the cingulate gyrus and cerebellum (Arsalidou & Taylor, 2011; Faye, 2019). Invasive modalities such as intracranial electroencephalography have also affirmed the TCM, contributing to a more robust understanding of the neuronal spatiotemporal dynamics during number processing (Daitch et al., 2016; Dastjerdi et al., 2013; Hermes et al., 2017; Shum et al., 2013).

To date, iEEG investigations have primarily relied on electrocorticography (ECoG) in the form of subdural strips and grids to analyze cortical areas. Despite compelling evidence that subcortical structures play an integral role in number processing (Chochon et al., 1999; Collins et al., 2017; Hofstetter & Dumoulin, 2021; Kutter et al., 2018), exceptionally few studies have conducted electrophysiological recordings from deeper subcortical regions (Kutter et al., 2018; Kutter et al., 2022). To address this gap, we leveraged the use of stereotactic electroencephalography (sEEG) electrodes in medically refractory epilepsy patients implanted for the purposes of seizure localization. Moreover, sEEG provides extensive coverage of brain areas implicated in subserving number processing as posited by the TCM. In this study, we sought to investigate the utility of sEEG recordings as a means of analyzing neural correlates of human number cognition. Drawing upon the TCM’s postulates, we presented 13 subjects with a number recognition task of numerical stimuli that differed at the level of representation type (symbolic vs. non-symbolic) and mode of stimuli delivery (visual vs. auditory). Using a classification algorithm and stimuli evoked sEEG recordings, we examined the extent to which number encoding brain regions aligned to the TCM. Additionally, this approach enabled us to explore neural substrates outside of this predicted framework.

## MATERIAL AND METHODS

### Participants

Thirteen patients with a diagnosis of medically refractory epilepsy were implanted with sEEG electrodes for the purpose of seizure onset zone localization. The sEEG electrodes used were 0.8 mm in diameter with a center-to-center pitch of 3.3 mm to 5mm between electrode contacts (PMT, Chanhassen, MN, USA). A total of 2,482 contacts were analyzed and distributed within each subject as shown in Figure 1. Pertinent demographic characteristics are tabulated in Table 1. Institutional Review Board approval was obtained at Oregon Health & Science University. All participants were over the age of 18 and provided written informed consent. Informed consent was obtained under the Declaration of the Principles of Helsinki.

**Table 1.**
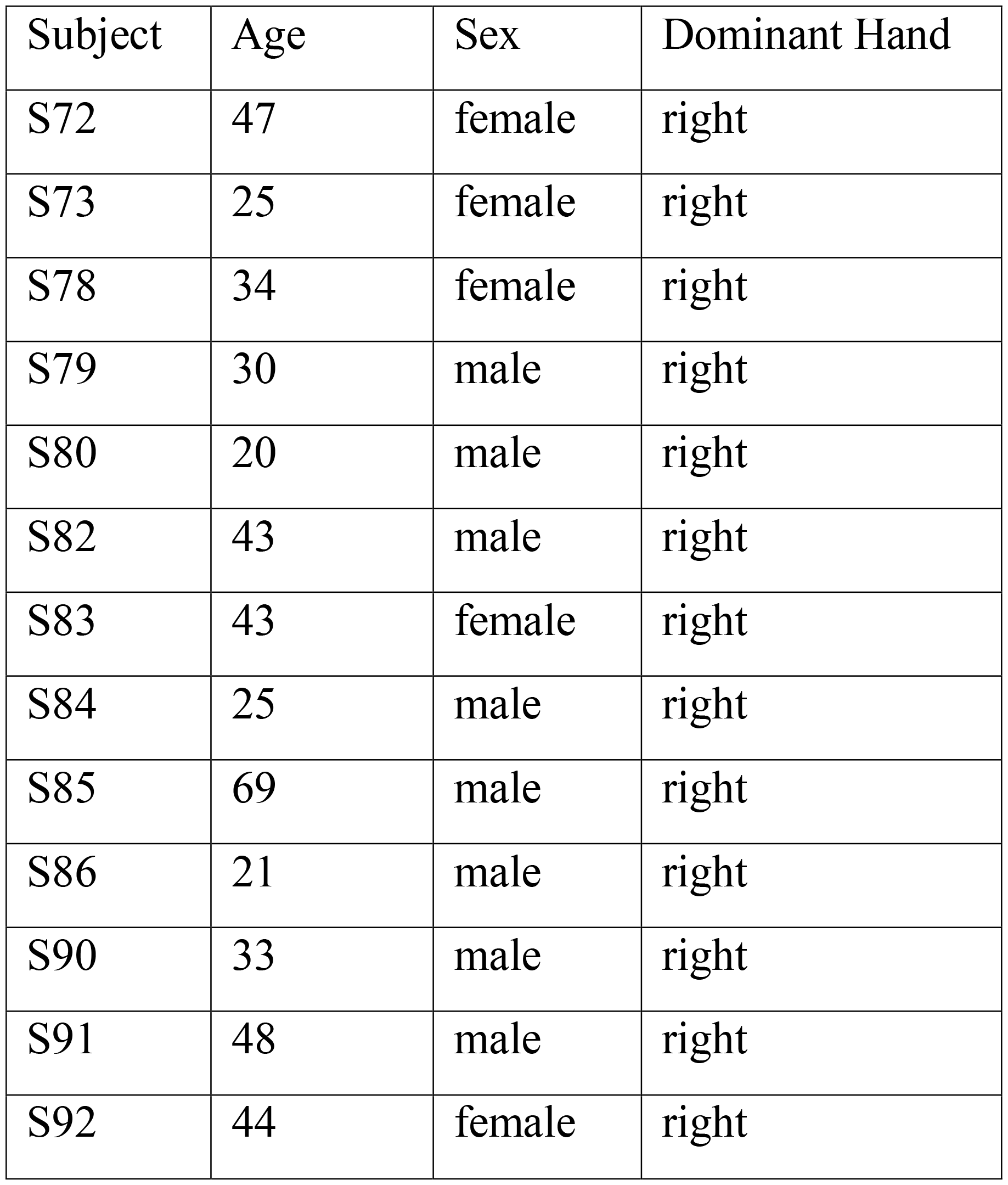
Subject demographics.

| Subject | Age | Sex | Dominant Hand |
| --- | --- | --- | --- |
| S72 | 47 | female | right |
| S73 | 25 | female | right |
| S78 | 34 | female | right |
| S79 | 30 | male | right |
| S80 | 20 | male | right |
| S82 | 43 | male | right |
| S83 | 43 | female | right |
| S84 | 25 | male | right |
| S85 | 69 | male | right |
| S86 | 21 | male | right |
| S90 | 33 | male | right |
| S91 | 48 | male | right |
| S92 | 44 | female | right |

**Fig 1.**
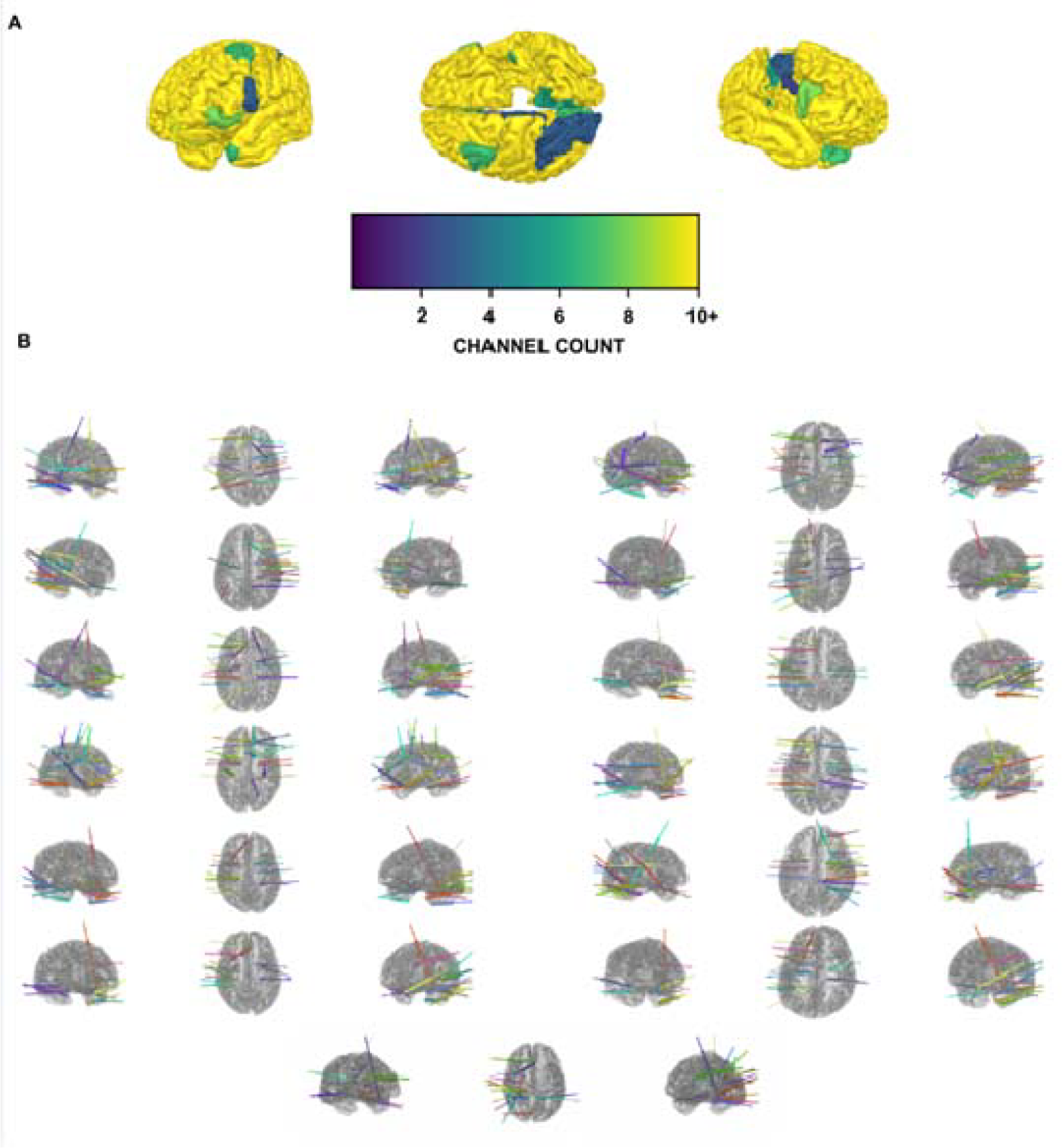
Electrode localization (A) Heat map of sEEG electrode contact distribution (B) sEEG implantation layouts for each of the 13 patients

### Behavioral Task

Patients performed a passive numerical recognition task on a laptop **(Figure 2)**. During the task, patients were presented with a number quantity from one to nine in one of four different representation formats. Two of the four were auditory (spoken number, sequential beeps) while the remaining two were visual (Arabic numeral, assortment of dots). These representation formats were designed to reflect common types of number stimuli encountered in daily life. The duration of each number trial was 1 second long with an intertrial period of 2 seconds. All 13 patients were presented with 199 number trials. To ensure attentiveness, patients were periodically presented with catch trials and prompted to press either left or right arrow keys which corresponded to whether the presented catch trial numerical quantity was odd or even, respectively. The task would not proceed until patients responded to the catch trial.

**Fig 2.**
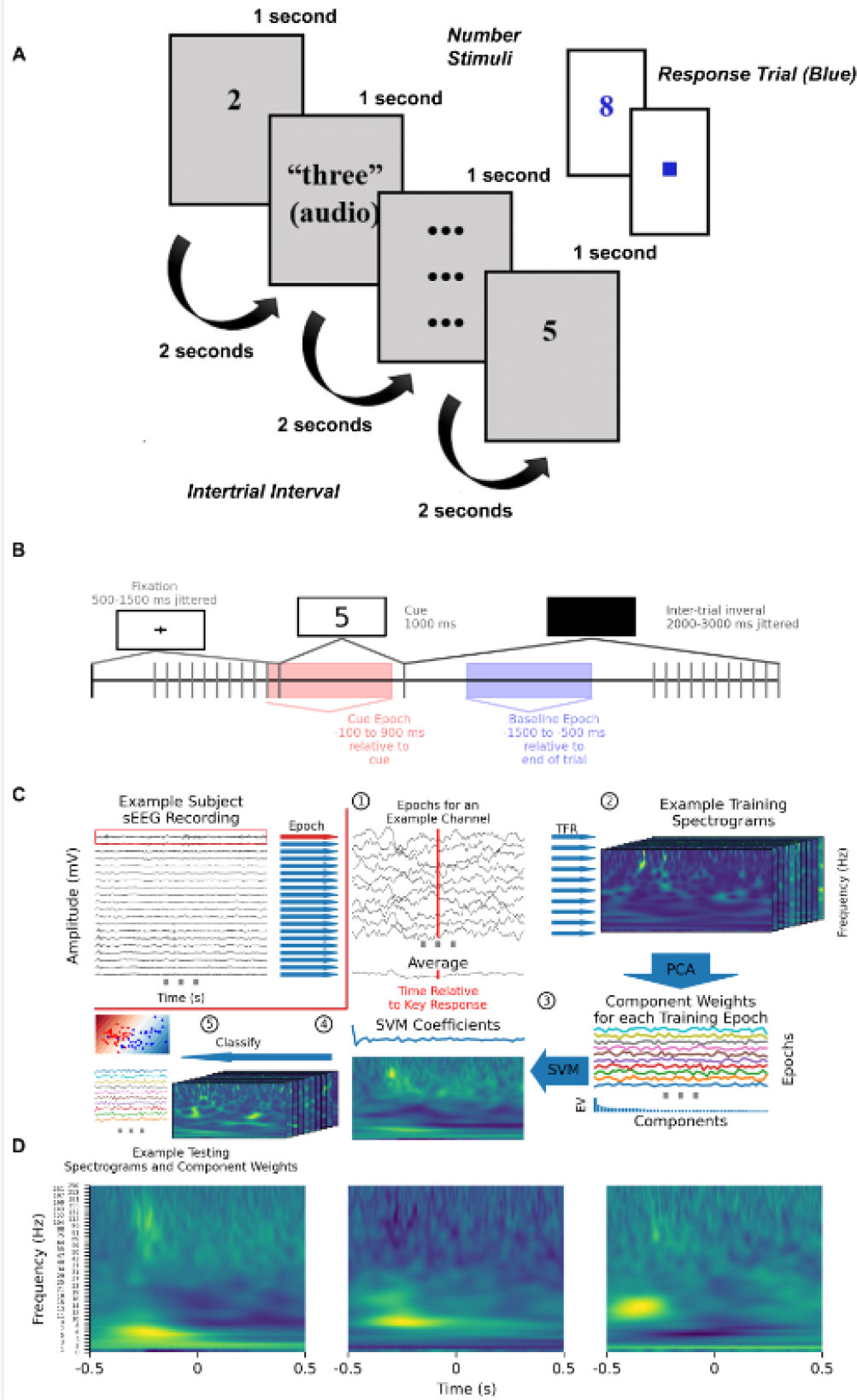
Task paradigm and classification algorithm: (A) Numerical task paradigm (B) Spliced epochs used in the classification algorithm (C) Schematic of the classification analysis pipeline using an exemplar contact. This process is iteratively performed for all sEEG contacts. Epochs are spliced (1) and decomposed into spectrograms (2). A PCA is then applied to the spectrograms for training trials (3). For components, the explained variance is shown as a bar plot beneath PCA weights per component. A SVM is fit to the component weights, multiplied by the principal components, summed, and projected back into spectrograms (4). Finally, coefficients are multipled and summed with the component weights from each test trial and classified based on the category of numerical representation format. The linear decision boundary for the first 2 principal components is shown as an example (5). (D) For this exemplar contact, the first three eigenvector spectrograms from the PCA decomposition are shown.

The task was administered using a custom jsPsych script implemented through a web browser. The laptop was placed in front of patients on a table comfortably positioned over their lap. Trials were synchronized to intracranial electrophysiology using a photodiode connected into the same amplifier as the sEEG data and attached to either the right or left corners of the laptop screen (Rockhill, 2020). Trials during corrupted photodiode events were excluded as accurate timing of intracranial electrophysiology changes could not be ascertained without photodiode synchronization. All participants included were right-hand dominant and responded to catch trials accordingly.

### Electrode localization

To determine sEEG electrode positions, preoperative stereotactic T1 and T2 magnetic resonance (MR) were registered to postoperative computerized tomography (CT) imaging studies with MNE-python (Rockhill, 2022). Anatomic labels were assigned to contacts using the Desikan-Killiany atlas label of each patient’s Freesurfer reconstruction (Fischl, 2012). Contact locations were warped to a template brain (cvs_avg35_inMNI152) to standardize contact positions across patients and allow between patient comparisons. Task-related cue and response events were synchronized using differences in time stamps recorded by the task computer relative to the time that the fixation stimulus was displayed which was synchronized by the photodiode. Using MNE-python, time-frequency spectrograms for each event were computed using the Morlet wavelets method with frequencies from 1 to 250 Hz. After being bandpass filtered between 0.1 and 40 Hz, voltage time-series signals were appended to the bottom of each spectrogram to account for event-related potentials in our classification analysis.

### Classification

We leveraged a classification analysis similar to Rockhill et al. 2022 to determine the classification accuracy of sEEG contacts, parcellated brain regions, and spectral features for all numerical stimuli (**Figure 2**). Number trial spectrograms consisted of the period -0.1s prior to number stimulus presentation to 0.9s afterwards. These spectrograms were classified differently from inter-trial interval spectrograms of the same duration. Training spectrograms were dimensionally reduced with principal component analysis (PCA). The first 50 principal components were used as inputs into a linear support vector machine (SVM) classifier deployed with scikit-learn (Buitinck, 2013). A 70/30% training-test split was used. A separate portion of the inter-trial interval was used as a null classification. Coefficient matrices from the SVM were validated using a one sample cluster permutation test with a significance threshold set at 99% of a T-distribution (alpha = 0.01) and degrees of freedom set at 12 (one less than the number of subjects). Within sEEG channels, clusters were deemed statistically significant if their T-statistics was greater than 99% of permuted clusters.

## RESULTS

### SVM classifier accuracy

Our linear SVM successfully classified spectrograms during number stimuli and specific representation formats from those during the intertrial interval. With an alpha threshold of 0.01 relative to the null distribution, we identified contacts with statistically significant classification probabilities. When all number stimuli were classified against the intertrial period, 429 had significant classification probabilities. For our four representation formats, 407, 776, 854, and 513 contacts had significant classification values during Arabic, beep, spoken, and dots, respectively (**Figure 3**).

**Fig 3.**
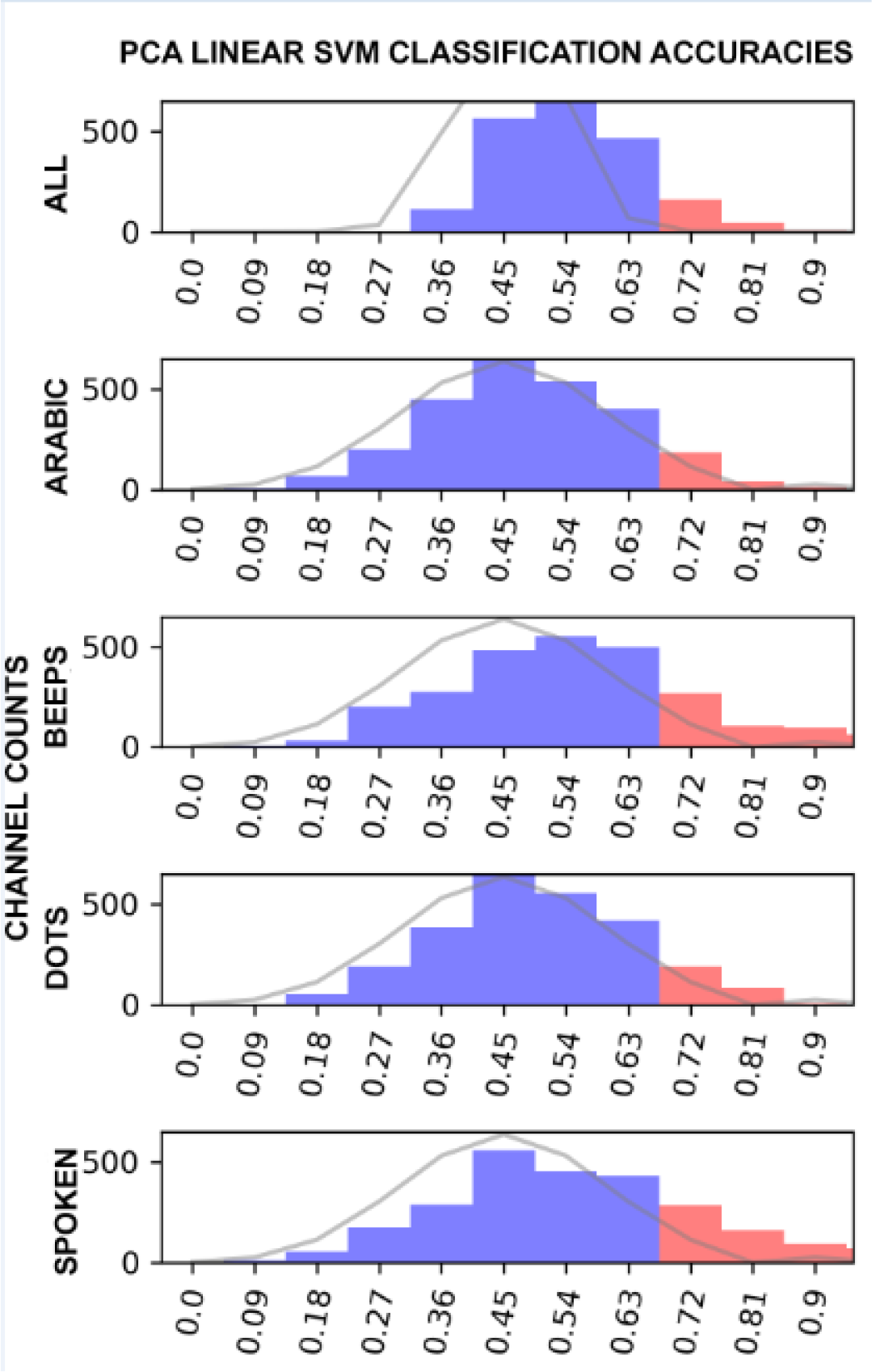
Histograms presenting the counts of sEEG channels with significant (red) and nonsignificant (blue) classification accuracy at alpha = 0.01 relative to a null distribution (grey line) across all contacts (paired t-test comparing difference in means between test and null distribution, p<0.00001). The distribution of significant and nonsignificant sEEG channels are presented for each number representation format.

### All Number Formats vs. Intertrial Period

We began by classifying spectrograms during all number stimuli, irrespective of representation format, from spectrograms during the intertrial interval. The SVM classification demonstrated that the bilateral parietal regions and the right lingual cortex had the highest classification accuracy (**Figure 4A**). Closer examination of the subcortex revealed that the right and left putamen had the highest classification accuracy of all subcortical structures with the right having a higher classification value than the left (**Figure 4A**). The three contacts with the highest classification value across all number stimuli are presented on Figure 4B. The first was in the right lingual cortex. Decreases in gamma, beta, and low frequency band activity before increases in the beta band classified numerical stimuli. The second was in the left inferior parietal cortex. Synchronized gamma, low beta, and theta band classified number stimuli presentation. The third was in the middle temporal cortex where theta and gamma decreases and increases in the alpha and beta frequencies were used to classify number stimuli.

**Fig. 4.**
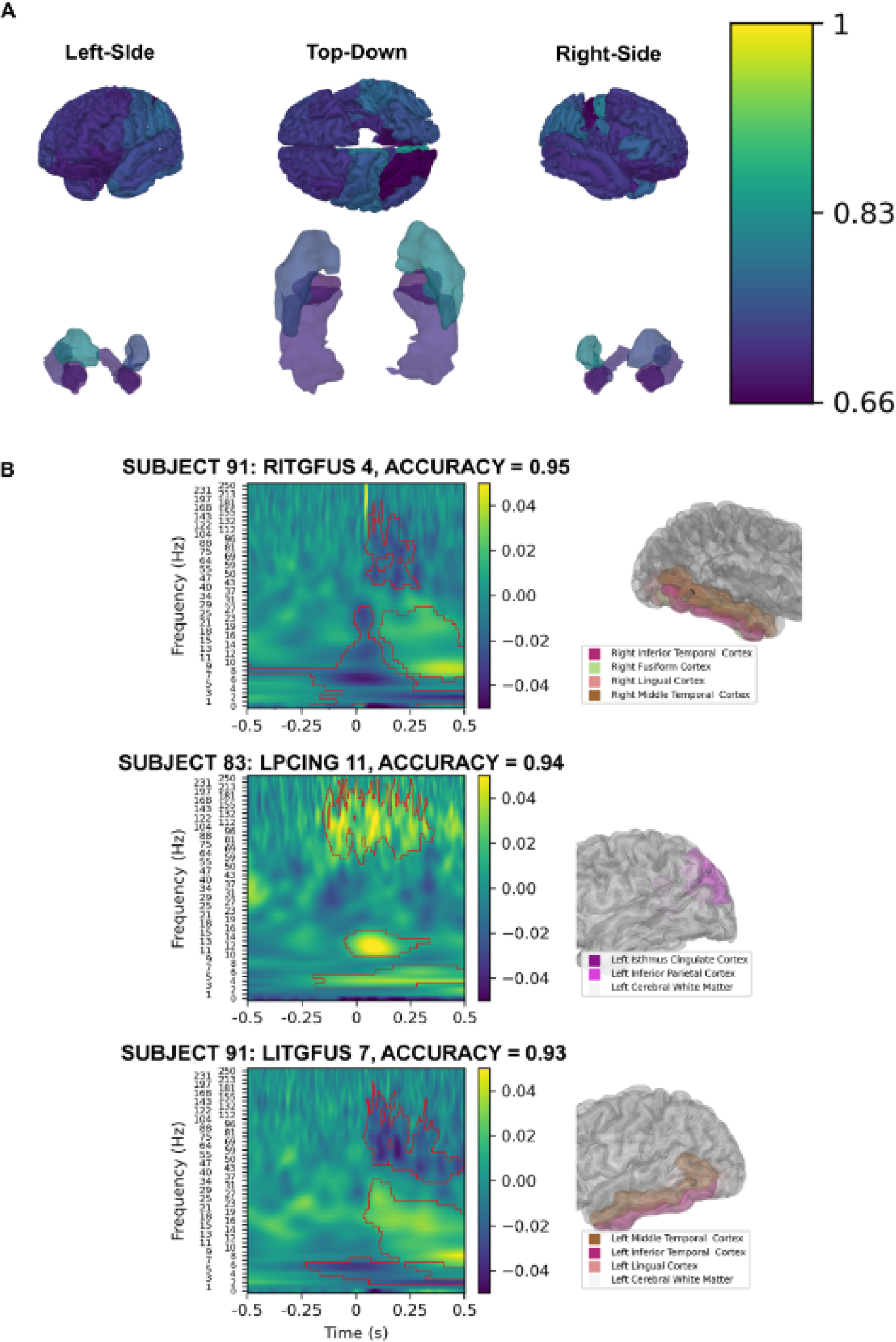
Linear SVM classification of spectrograms from all number trials versus intertrial interval spectrograms. Classification value of parcellated cortical (top row) and subcortical (bottom row) brain regions are presented with a gradient (Yellow; 1 = 100% classification accuracy, Purple: 0.66 = 66% classification accuracy) (A). The three contacts with the highest classification value are shown in (B). Left panels show SVM coefficients from spectrogram classification in the red contours. Right panels show the location of the contact within parcellated brain regions.

### Classification of Auditory Representation Formats

SVM classification of beep spectrograms showed that bilateral superior temporal and inferior parietal cortex had robust classification values **(Figure 5A)**. Both right and left putamen had strong classification value though the right putamen was higher (**Figure 5A**). Of the three contacts with the highest classification accuracy, two were in the left superior temporal gyrus while the third was in the right superior temporal gyrus (**Figure 5B**). Gamma and theta to low beta increases were used for classifying beeps in the right superior temporal gyrus contact. These changes were also present and used for classification in one of the two left superior temporal gyrus contacts while theta to alpha increases were used for classification in the other superior temporal gyrus contact.

**Fig. 5.**
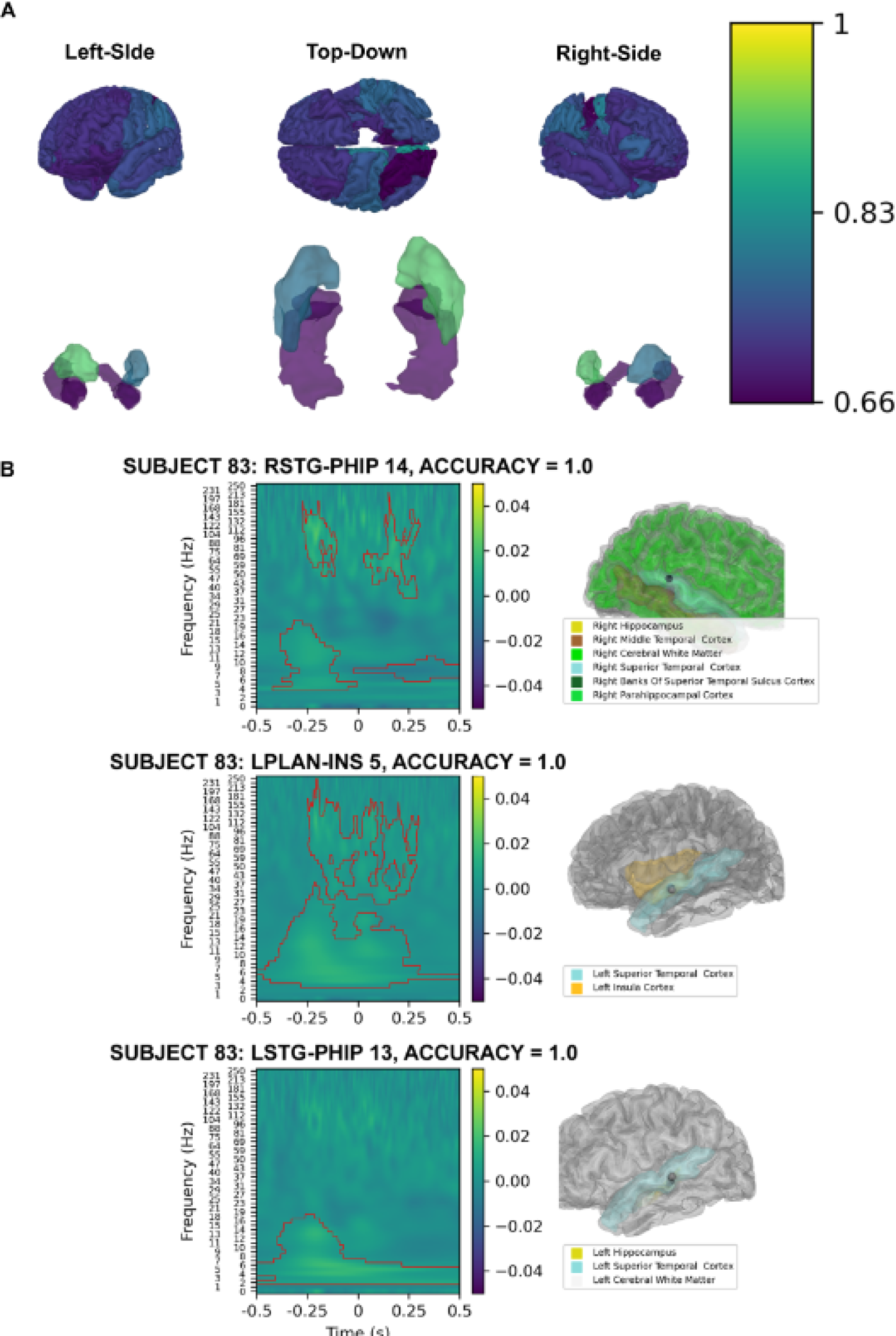
Linear SVM classification of spectrograms from sequential beep trials versus intertrial interval spectrograms. Classification value of parcellated cortical (top row) and subcortical (bottom row) brain regions are presented with a gradient (Yellow; 1 = 100% classification accuracy, Purple: 0.66 = 66% classification accuracy) (A). The three contacts with the highest classification value are shown in (B). Left panels show SVM coefficients from spectrogram classification in the red contours. Right panels show the location of the contact within parcellated brain regions.

Similar to the classification of beeps, spoken number trials demonstrated the best classification value in the bilateral superior temporal cortices (**Figure 6A**). Within the subcortex, the bilateral putamen demonstrated robust classification value similar to that of the superior temporal cortices (**Figure 6A**). The three contacts of highest accuracy were found in the bilateral superior temporal cortices and the right supramarginal cortex (**Figure 6B**). A decrease in the theta to alpha frequency bands was used to classify spoken number trials from intertrial intervals within the right supramarginal gyrus. High gamma increases were used to classify spoken number trials within both superior temporal cortices. Additionally, an initial theta increase followed by theta/alpha increases were used within both superior temporal cortex contacts for classification as well.

**Fig. 6.**
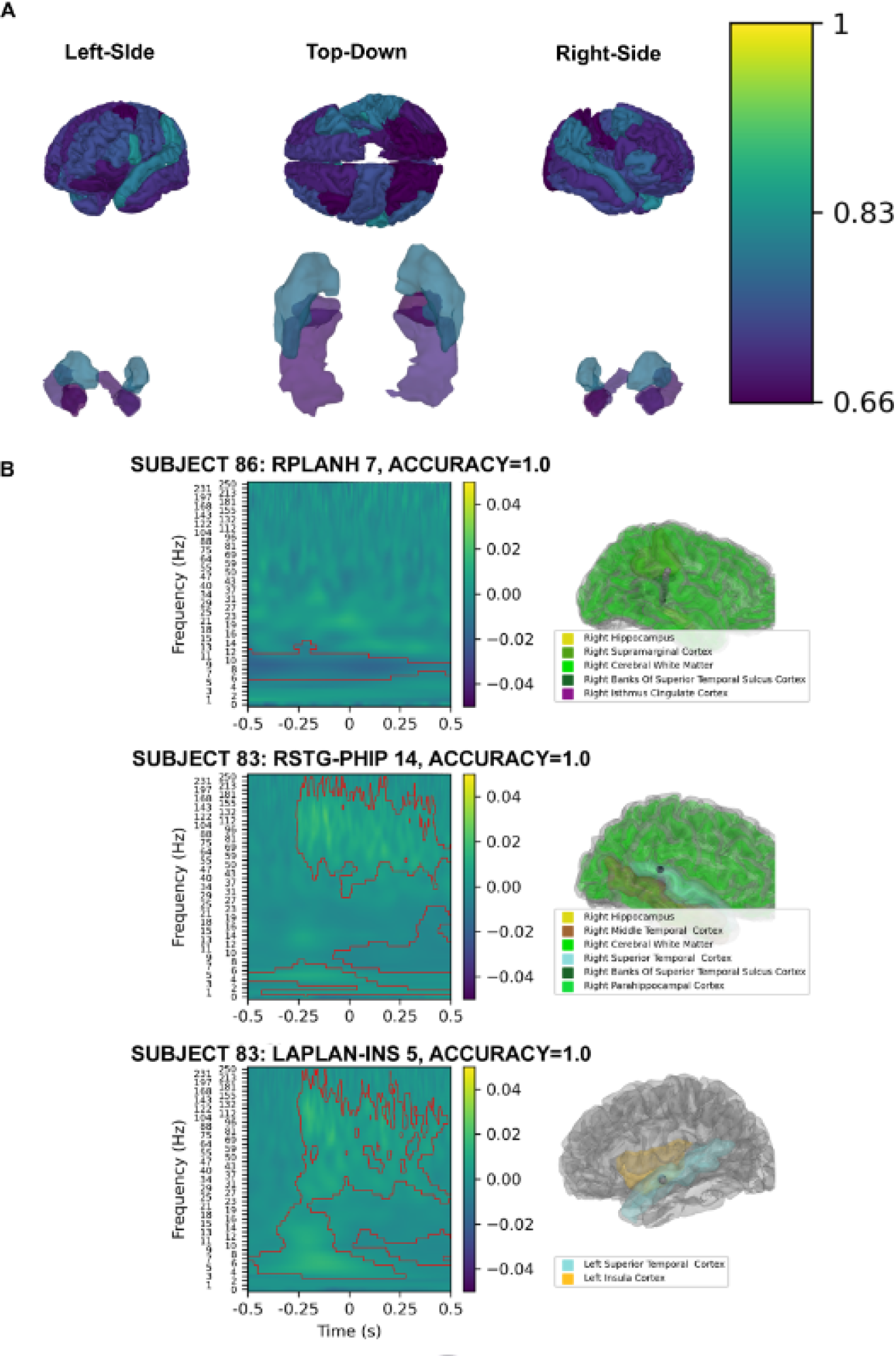
Linear SVM classification of spectrograms from all spoken number trials versus intertrial interval spectrograms. Classification value of parcellated cortical (top row) and subcortical (bottom row) brain regions are presented with a gradient (Yellow; 1 = 100% classification accuracy, Purple: 0.66 = 66% classification accuracy) (A). The three contacts with the highest classification value are shown in (B). Left panels show SVM coefficients from spectrogram classification in the red contours. Right panels show the location of the contact within parcellated brain regions.

### Classification of Visual Representation Formats

The SVM classification of spectrograms during Arabic numeral trials indicated that the frontal lobes and parietal lobe regions posterior to the postcentral gyrus had robust classification value (**Figure 7A**). Of note, the inferior precentral gyrus had the best classification accuracy, however, this region had a sparse number of contacts. The right putamen had a particularly robust classification value, superseding that of all parcellated cortical and subcortical structures (**Figure 7A**). The three contacts of highest classification accuracy were localized in the supramarginal, middle temporal cortex, and left insula (**Figure 7B**). Decreases and increases in the theta to beta frequency range were used to classify Arabic numerals for the contacts located in the supramarginal and middle temporal cortex, respectively.

**Fig. 7.**
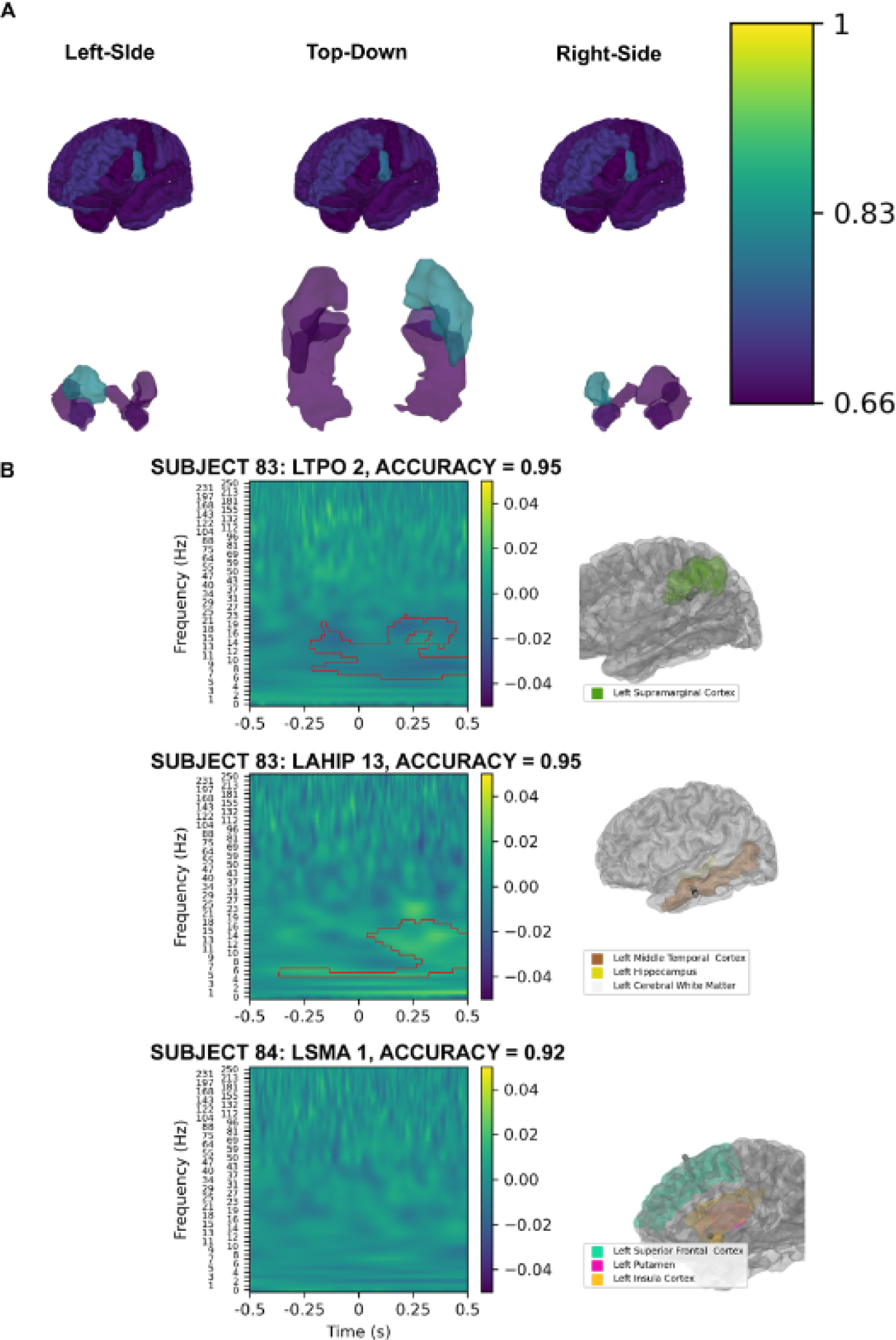
Linear SVM classification of spectrograms from all Arabic numeral trials versus intertrial interval spectrograms. Classification value of parcellated cortical (top row) and subcortical (bottom row) brain regions are presented with a gradient (Yellow; 1 = 100% classification accuracy, Purple: 0.66 = 66% classification accuracy) (A). The three contacts with the highest classification value are shown in (B). Left panels show SVM coefficients from spectrogram classification in the red contours. Right panels show the location of the contact within parcellated brain regions.

Despite having fewer contacts than other regions, the left parietal cortex had the best classification value when the SVM classified dot spectrograms from intertrial spectrograms (**Figure 8A**). The left frontal lobe also had robust classification value as well. Within the subcortex, the right putamen had robust classification accuracy, superseding that of other subcortical regions and most cortical regions besides the left parietal cortex (**Figure 8A**). The three contacts with the highest classification accuracies were in the left fusiform, right fusiform, and right superior frontal cortex (**Figure 8B**). A beta increase was used to classify dot trials in the right fusiform while a theta to beta frequency band increase was used for classification in the right superior frontal cortex.

**Fig. 8.**
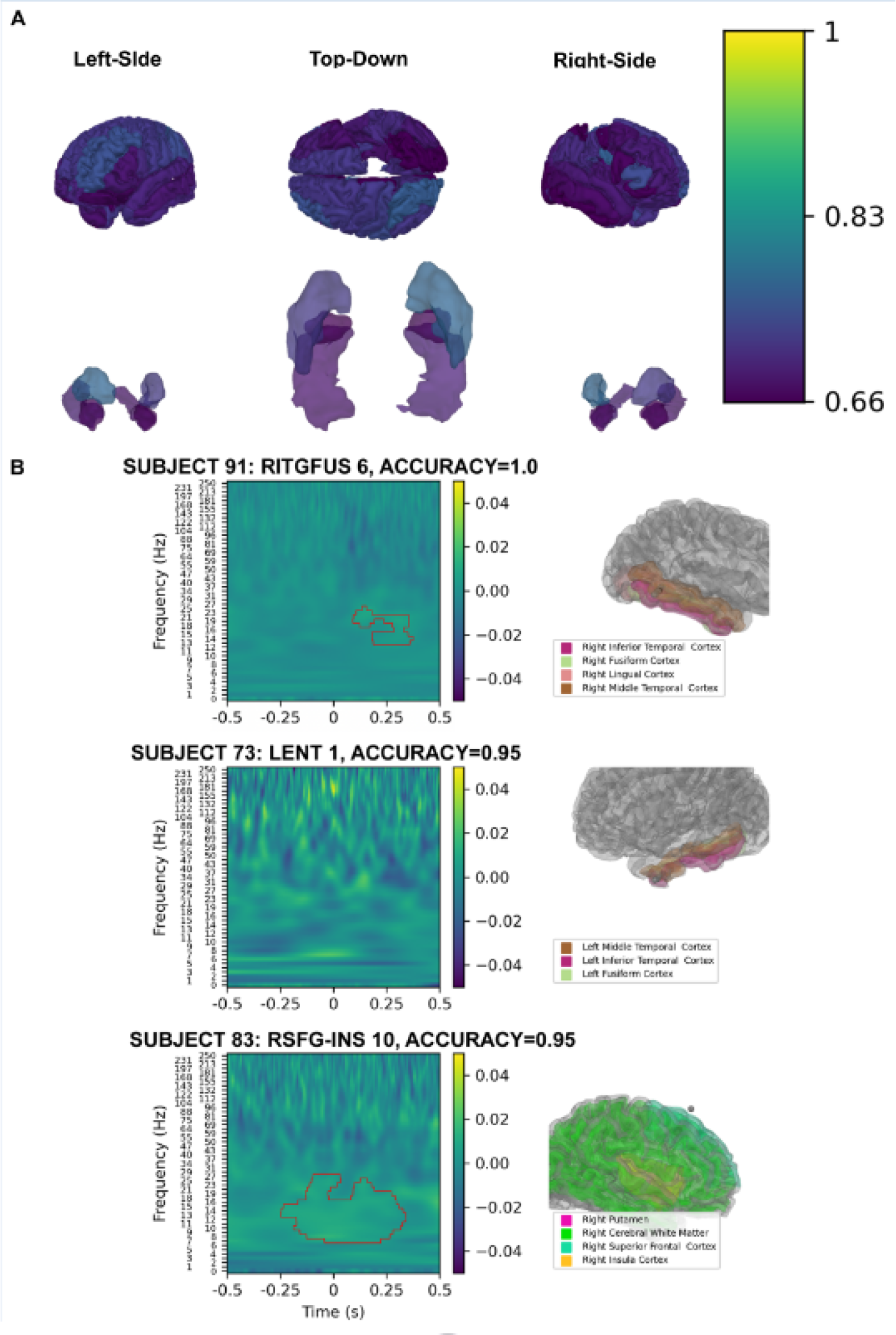
Linear SVM classification of spectrograms from all assorted dots trials versus intertrial interval spectrograms. Classification value of parcellated cortical (top row) and subcortical (bottom row) brain regions are presented with a gradient (Yellow; 1 = 100% classification accuracy, Purple: 0.66 = 66% classification accuracy) (A). The three contacts with the highest classification value are shown in (B). Left panels show SVM coefficients from spectrogram classification in the red contours. Right panels show the location of the contact within parcellated brain regions.

### Spectral Features with High Classification Value for Number Stimuli

To better understand the SVM classification’s coefficient matrices, we generated feature maps showcasing the relative abundance of statistically significant time-frequency clusters and the proportion of positive significant clusters to determine the directional patterns of time-frequency cluster changes (**Figure 9**). Then, we assessed the classification value of these cluster changes. Across all number representation formats, significant clusters within the theta to beta frequency ranges were well represented. To a less robust degree, spoken number and beeps demonstrated a number of significant gamma band clusters as well. The theta and beta clusters generally increased in power for all representation formats with the exception of spoken number, where beta clusters demonstrated an initial increase before a decrease. All representation formats, specifically beeps and spoken number with significant gamma band clusters had an increase in gamma power associated with these significant gamma clusters. Increases in gamma had the highest classification accuracy. Meanwhile alterations in the alpha and beta frequency bands had less robust but greater than chance classification accuracy.

**Fig. 9.**
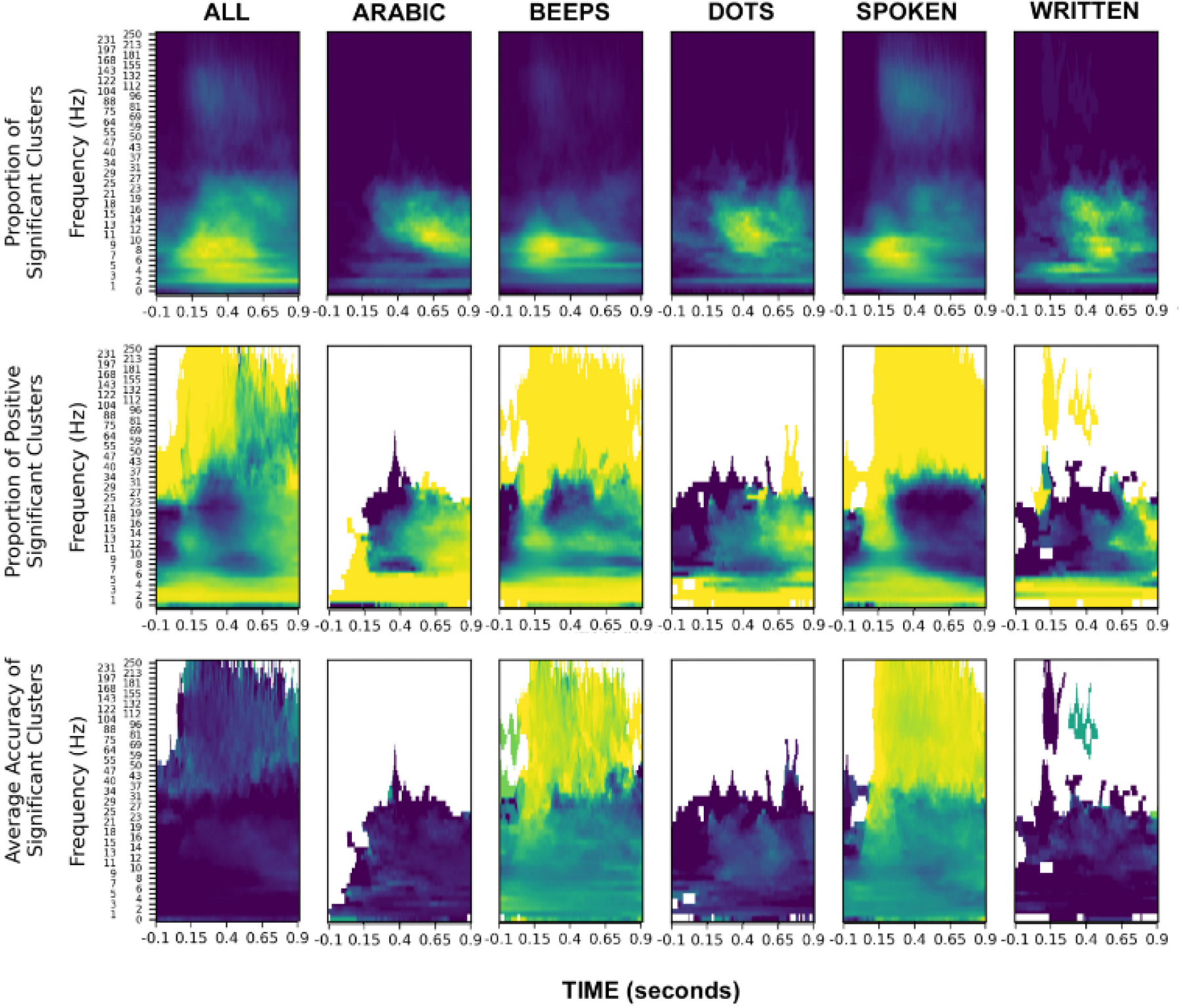
This is a summary map of all contacts with statistically significant SVM classifications. The top row presents the proportion of significant SVM clusters. Yellow indicates a greater proportion of clusters while blue represents few clusters at specific time-frequency points. The middle row shows the directionality of spectral change for these significant SVM clusters during number trial conditions versus intertrial intervals. Yellow indicates an increase in time-frequency clusters during number trials versus intertrial intervals while blue indicates a decrease in time-frequency clusters. White regions in the plot indicate that no directional change occurred. The bottom row presents the classification value of the directionality of SVM cluster changes. Yellow reflects high classification accuracy close while dark blue represents less robust classification accuracy. Time-frequency points with no significant classification value are white.

## DISCUSSION

Using a linear SVM classifier with PCA, we identified cortical and subcortical structures, individual contacts, and spectral patterns with high classification value during our number stimuli trials. Overall, our findings align with results from prior fMRI and ECoG investigations into number cognition (Arsalidou & Taylor, 2011; Daitch, 2016; Faye, 2019; Shum, 2013), and by extension, lie congruent with existing neuroanatomical models of number cognition.

When number stimuli, irrespective of representation format, were classified against the intertrial period, we found evidence of bilateral parietal lobe engagement and robust engagement of the left inferior parietal cortex, which aligns with the postulates of the TCM (Dehaene & Cohen, 1995). It has been hypothesized that the intraparietal sulcus is responsible for encoding an innate, conserved sense of nonabstract number processing (Arsalidou & Taylor, 2011; Dehaene & Cohen, 1995). Previous fMRI studies have implicated the involvement of this region across a variety of number stimuli which has been redemonstrated in ECoG studies (Dastjerdi, 2013; Vogel, 2017; Zago, 2008). We also observed that auditory numerical stimuli reliably engaged their putative cortical substrates. For both beeps and spoken number trials, the bilateral superior temporal cortices had good classification value, suggesting preferential activation at these sites. This is congruent with known localizations of the eloquent human auditory cortex which have been implicated by fMRI analyses to underline auditory number processing (Chee, 1999; Howard, 2000; Piazza, 2006).

Classification of the visual representation formats did not consistently implicate the expected cortical regions of engagement. Prior ECoG analyses have identified a visual number form area (VNFA) within the fusiform and inferior temporal gyri that is preferentially engaged in response to Arabic numerals (Hermes, 2017; Shum, 2013). Instead, during Arabic trials, we found evidence of frontal lobe involvement with evidence of inferior parietal and middle temporal cortical engagement. It is likely that our sEEG contacts did not adequately sample from the VNFA, biasing the classification results towards sites with better contact coverage. Finally, classification of dot trials indicated preferential engagement of the left parietal cortex, as expected, as well as the left frontal lobe. This aligns with the frontoparietal network of number cognition posited by Dehaene et al. and the hypothesized role of the left parietal rgcortex put forth by the TCM (Dehaene & Cohen, 1995; Dehaene, 2003).

Interestingly, across all number representation formats, the right putamen had the best classification value of all subcortical structures. It also demonstrated classification accuracy comparable or superior to that of putative cortical substrates of number processing. It is unclear to why the right putamen had consistently superior classification to the left putamen. However, this may be partially related to the hemispheric dominance of our patients as all were left hemisphere dominant. Although commonly associated with motor function as a component of the basal ganglia, the putamen is known to take part in higher-level cognitive functions as well (Ketteler, 2008; Mestres-Missé, 2010; Viñas-Guasch & Wu, 2017). Enhanced putaminal engagement during numerical processing has been demonstrated during magnitude evaluation and arithmetic tasks with functional imaging (Chochon, 1999; Hofstetter & Dumoulin, 2021). Recently, using a high field MRI, investigators recently detected tuned neural responses to numerical quantities within the putamen during a tactile numerosity task (Hofstetter & Dumoulin, 2021), illustrating the pertinent role of the putamen in integrating and comprehending numerosity inputs.

Considering that neuroimaging studies have also demonstrated the putamen’s role in language cognition (Ketteler, 2008; Mestres-Missé, 2010), the engagement of the putamen during our number recognition paradigm touches on the potential interplay between number cognition and language. We did not attempt to compare neuroanatomic and electrophysiologic features of number stimuli to language ones in our study due to time constraints. There is spirited debate, particularly within neuropsychology and social sciences literature, over whether the development of and capacity for numerical cognition is independent of language acquisition (Gelman & Butterworth, 2005). More recent evidence suggests that the development of large and exact numerosity representations is contingent upon access to language (Spaepen, 2011). As a corollary, conceptualizing smaller and less precise quantities may be agnostic of language, which suggests that there are indeed numerosity specific, or language independent, neural substrates circuitry. Disentangling numerical circuits from language ones could be of clinical value, especially in the setting of Gerstmann syndrome which is classically typified by acalculia. To this end, future studies should leverage structural and functional connectivity methods while directing specific attention towards the involvement of subcortical structures.

Because fMRI analyses hinge upon blood-oxygen-level-dependent (BOLD) signal changes, which are strongly coupled with gamma frequency local field potentials (Crone, 2001), prior iEEG studies into number cognition have largely directed their attention to gamma band perturbations as a marker for synchronized neuronal activity (Daitch, 2016; Dastjerdi, 2013; Hermes, 2017; Shum, 2013). While gamma band activity had strong classification value within our analysis, our time-frequency cluster feature maps suggest that power changes in lower frequency bands may also be of value in distinguishing numerical stimuli from non-numerical stimuli and characterizing specific representation formats. Future studies should attempt to further elucidate the role of theta through beta frequency band activity during number processing.

There were several limitations to our study. First, by only using sEEG, the spatial resolution of our cortical recordings was constrained, as placement of electrodes are solely determined by clinical purposes. Nonetheless, we were still able to associate number stimuli with structures that align with existing models of number cognition, specifically the TCM. Second, we chose to use a linear SVM with PCA as our classification method knowing that this may come at the sacrifice of classification accuracy. We opted against a more complex and robust classification method with the intent of prioritizing interpretability ahead of classification accuracy. Despite these limitations, to our knowledge, this is the first study to utilize sEEG depth electrodes for the purpose of investigating human number cognition through sampling both cortical and subcortical structures.

In conclusion, we used a machine learning classifier to identify cortical and subcortical substrates of human number cognition. Our findings support postulates of the TCM in number processing. However, we also determined that subcortical structures, particularly the putamen exhibited robust classification accuracy in response to numerical stimuli, thus expanding this framework. Analyses of spectral feature maps revealed that non-gamma frequency bands held greater than chance classification value and could be potentially used to characterize format specific number representations. We provide both neuroanatomical and electrophysiologic targets of interest that can be leveraged in future number cognition investigations.

## AUTHOR CONTRIBUTIONS

H.T.: data curation, investigation, project administration, writing—original draft, writing— review and editing; A.P.R: data curation, methodology, software, writing—review and editing; C.L.R: data curation, project administration, writing—review and editing, C.N.: data curation, investigation, project administration, B.S.: writing—review and editing, investigation, A.F.: data curation, writing—review and editing, investigation; M.N.S.: data curation, writing—review and editing, investigation; M.I.: writing—review and editing, investigation, D.R. C.: writing—review and editing, K.L.C: resources, supervision, writing—review and editing, A.M.R: conceptualization, investigation, project administration, resources, supervision, writing—review and editing.

